# Gap junction mediated feed-forward inhibition ensures ultra precise temporal patterning in vocal behavior

**DOI:** 10.1101/2020.05.27.119339

**Authors:** Boris P. Chagnaud, Jonathan Perelmuter, Paul Forlano, Andrew H. Bass

## Abstract

Precise neuronal firing is especially important for behaviors highly dependent on the correct sequencing and timing of muscle activity patterns, such as acoustic signalling. We show that extreme temporal precision in motoneuronal firing within a hindbrain network that directly determines call duration, pulse repetition rate and fundamental frequency in a teleost fish, the Gulf toadfish, depends on gap junction-mediated, feed-forward glycinergic inhibition that generates a period of reduced probability of motoneuron activation. Super-resolution microscopy confirms glycinergic release sites contacting motoneuron somata and dendrites. Synchronous motoneuron activity can also induce action potential firing in pre-motoneurons, a feature that could figure prominently into motor timing. Gap junction-mediated, feed-forward glycinergic inhibition provides a novel means for achieving temporal precision in the millisecond range for rapid modulation of an acoustic signal and perhaps other motor behaviors.

## Introduction

Complex behaviors often depend on temporally precise neuronal firing that coordinates network activity at brain levels ranging from cortical microcircuits to hindbrain pattern generators (Kros et al., 2017; Llinás, 2014; Sober et al., 2018). Mechanisms known to increase precision at single cell and network levels include feed-forward inhibition in audition (Grothe, 2003), recurrent inhibitory input in cortex (Kapfer et al., 2007), and neuronal synchrony in cortical and sensory neurons (Tiesinga & Sejnowski, 2001; Uhlhaas et al., 2010). Synchronous, concurrent activation of neurons is widely distributed in the brain (Llinás, 2014) and especially important for behaviors requiring both rapid and precise motoneuron activation such as electrogenesis in fishes (Bennett, 1971), and vocalization in fishes (Chagnaud et al., 2012) and tetrapods (Kwong-Brown et al., 2019; Mead et al., 2017).

Several mechanisms by themselves or in combination contribute to neuronal synchrony and temporal precision: coherent excitatory firing, electrotonic coupling, and (recurrent) inhibitory input (Kapfer et al., 2007; Singer, 1999; Uhlhaas & Singer, 2006). While coherent (i.e., phasic) input leads to neuronal coupling mainly by excitation, inhibition might be the predominant way to synchronize activity (Van Vreeswijk et al., 1994). Electrotonic coupling enhances synchrony by spreading voltage changes, for example during synaptic inputs, that lead to concomitant membrane potential changes within an interconnected population (Bennett & Zukin, 2004; Pereda, 2014).

A system exhibiting extreme levels of synchronous activity, making it ideal to study mechanisms underlying precise neuronal firing, is the hindbrain vocal network of toadfishes, a marine order of teleosts that depends on acoustic signalling for social interactions (Bass et al., 2015). Motoneurons within each midline-positioned vocal motor nucleus (VMN) innervate the ipsilateral superfast vocal muscle attached to the swim bladder via a vocal nerve formed by two occipital nerve roots comparable to hypoglossal nerve roots (Fig. 1a). Hundreds of motoneurons in each VMN fire in phase within 1.5 milliseconds at frequencies of up to 250 Hz for time periods ranging from hundreds of milliseconds to minutes. Each VMN is bilaterally innervated by adjacent pacemaker neurons (VPN, Fig. 1b, c) that provide coherent excitatory input and determine VMN firing rate that directly translates into pulse repetition rate or fundamental frequency (Bass & Baker, 1990; Chagnaud et al., 2011; Chagnaud & Bass, 2014). Both VPN and VMN receive input from a more rostral vocal pre-pacemaker nucleus (VPP, Fig. 1b) that encodes duration (Chagnaud et al., 2011; Chagnaud & Bass, 2014).

**Figure 1:**
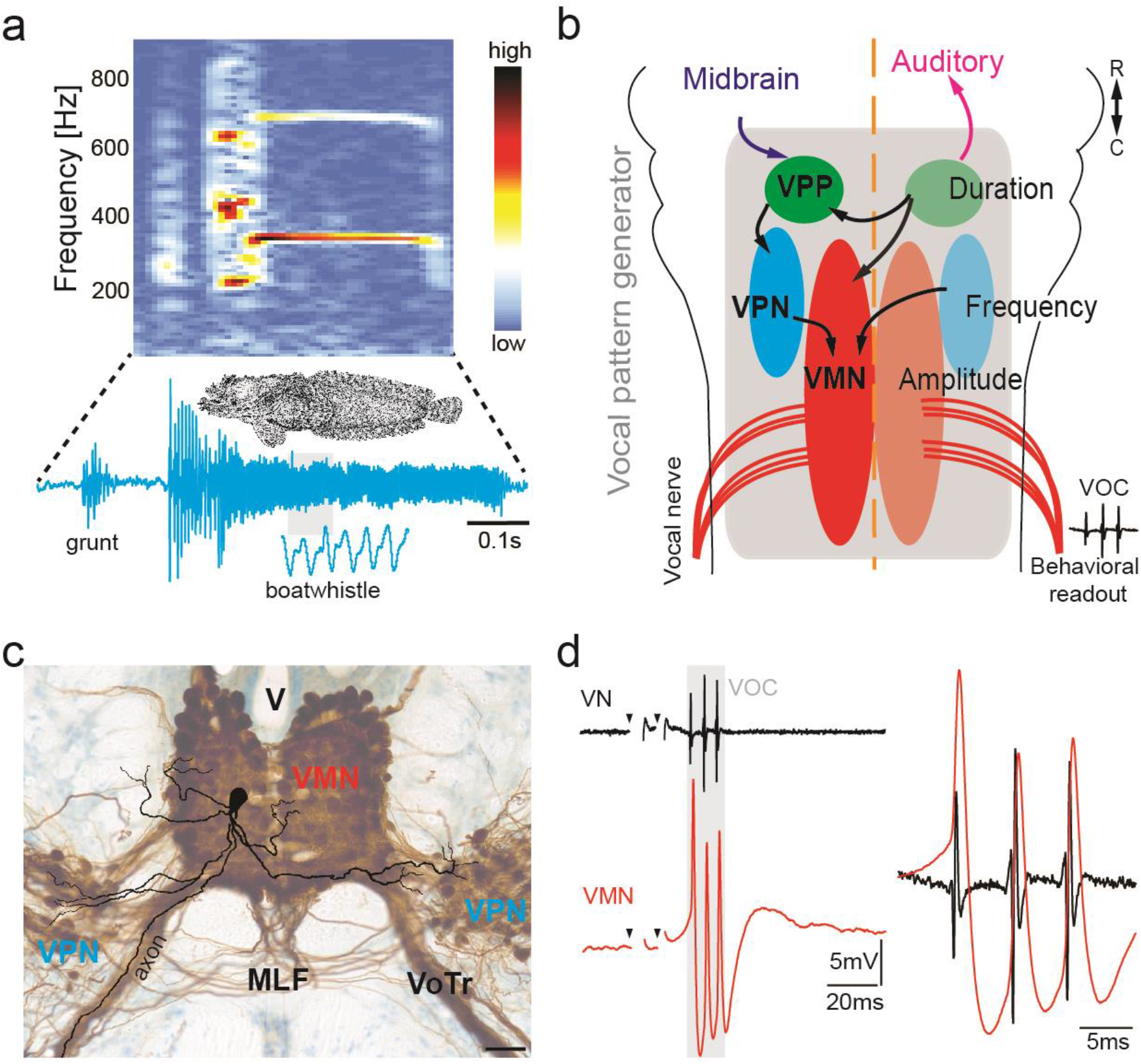
Toadfish vocalization and underlying vocal motor circuity. a) Spectrogram and waveform of a toadfish boatwhistle. b) Dorsal schematic view of caudal hindbrain showing toadfish vocal motor circuit comprised of three anatomically separate nuclei coding for different attributes: duration coding vocal prepacemaker nucleus (VPP), frequency (pulse repetition rate) coding vocal pacemaker nucleus (VPN) and amplitude coding vocal motor nucleus (VMN). Vocal nerves innervate muscles used in sound production. c) Photograph of transverse hindbrain section showing reconstructed neurobiotin-filled, VMN motoneuron superimposed over neurobiotin-filled VMN and VPN (dark brown); cresyl violet counterstain. d) Vocal nerve (VN) compound potential (fictive call VOC; grey box) and intracellular VMN motoneuron recording show highly time-locked activity. Scale bar in (c) represents 100 μm.

Earlier investigations of the VMN were among the first to show the potential contribution of electrotonic coupling to neuronal synchrony. This included ultrastructure evidence for gap junction contacts between motoneurons and axons of unidentified origin (likely VPN and VPP) (Bass & Marchaterre, 1989; Pappas & Bennett, 1966). More recent studies show that vocal nerve labelling with gap junction impassable tracers leads to dense retrograde labelling of the ipsilateral VMN, while gap junction passable tracers lead to dense transneuronal, bilateral labelling of VMN, VPN and VPP (Fig. 1c) (Bass et al., 1994; Chagnaud & Bass, 2014).

Here, we use intracellular, *in situ* recordings to investigate intrinsic and network properties underlying ultra-precise temporal firing of the VMN in Gulf toadfish (*Opsanus beta*, Fig. 1d). Like other toadfishes (e.g., see Rice & Bass, 2009), Gulf toadfish produce two main types of vocalizations, brief broadband agonistic grunts and multiharmonic advertisement calls known as boatwhistles (grunt pulse repetition rate and boatwhistle fundamental frequency are ~ 200-250 Hz; see Fig. 1d; also Bass et al., 2015; Maruska & Mensinger, 2009). Both sexes produce grunts, but only males are known to produce boatwhistles (Thorson & Fine, 2002; Winn, 1967; Winn, 1972). As shown, the VMN motoneurons display strong electrotonic coupling; coherent, high frequency (up to 250 Hz depending on temperature) excitatory input; and inhibitory glycinergic and GABAergic inputs, all known to contribute to coordinated network activity. We also show that vocal motoneurons exhibit a strong after-hyperpolarization (AHP) not seen during intracellular square pulse current injection that enhances synchronized activity in the vocal network. *In situ* pharmacology shows the AHP is mediated by a glycinergic, feed-forward inhibition dependent on gap junctional coupling; glycinergic release sites in contact with VMN somata and dendrites are confirmed with super-resolution microscopy. Synchronous motoneuron activity can also generate feedback activation of glycinergic VPN neurons. We propose that a gap junction-mediated, feed-forward glycinergic inhibition provides a novel means to enhance temporal precision in the activation of neural networks underlying acoustic signalling, and perhaps time coding in other motor systems.

## Results

### Vocal Nerve Physiology: Fictive Vocalization

Intracellular VMN motoneuron recordings were done *in situ*. Muscle activity was pharmacologically blocked and hindbrain vocal output monitored as compound potentials recorded intra-cranially from ventral occipital nerve roots (Fig. 1b, d). We refer to the highly-stereotyped pattern of nerve potentials that occur in the absence of vocal muscle activity as a fictive vocalization (VOC). Each VOC potential directly reflects synchronous VMN activity (Chagnaud et al., 2012). The rate (or frequency) and duration of VOC potentials directly determine the rate and duration of vocal muscle contractions that set, in turn, call pulse repetition rate and duration, respectively (Fig. 1a) (Bass & Baker, 1990; Cohen & Winn, 1967; Elemans et al., 2014; Pappas & Bennett, 1966; Remage-Healey & Bass, 2006; Rubow & Bass, 2009; Skoglund, 1961).

VOCs occurred spontaneously or were evoked by brief trains of low amplitude, electrical microstimulation in midbrain sites in a region comparable to the periaqueductal grey of birds and mammals (Kittelberger & Bass, 2013). Like other toadfishes, both vocal nerves fired in phase and individual VOC potentials were matched 1:1 with the activity of individual VMN motoneurons (Fig. 1d) (Bass & Baker, 1990; Chagnaud & Bass, 2014). Motoneurons had three to five main dendritic branches and an axon arising from a primary dendrite or soma that lacked axon collaterals and exited the brain ipsilaterally via the vocal tract (VoTr, Fig. 1c) (Bass & Baker, 1990; Chagnaud & Bass, 2014).

### Midbrain-evoked vocal motoneuron physiology

Electrical midbrain stimulation led to membrane depolarizations in motoneurons that increased in amplitude until a single action potential (AP) was fired (Fig. 2a_1,2_). This AP coincided with the presence of a single VOC nerve potential. With increasing stimulus strength, additional APs were detected that matched 1:1 with additional VOC potentials (Fig. 2a_3,4,5_) that together mimicked the pulse repetition rate of natural grunts (Fig. 1a) (Elemans et al., 2014; Maruska & Mensinger, 2009; Tavolga, 1958; Winn, 1967).

**Figure 2:**
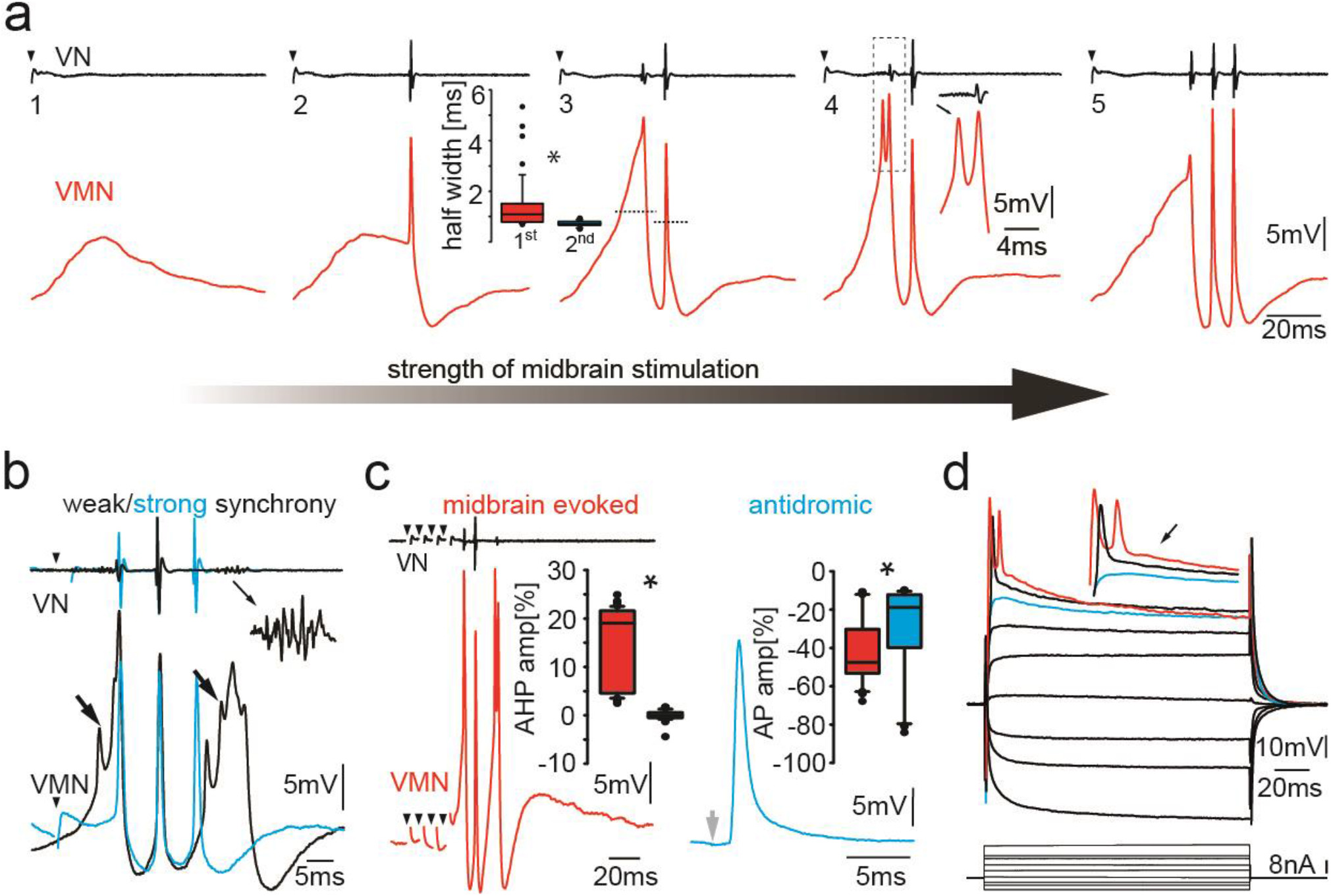
Vocal motor activity in motoneurons. a) Intracellular recordings of a single VMN motoneuron showing different stages of activity in the generation of vocal nerve (VN) motor volley, the fictive vocalization, elicited by midbrain electrical stimulation. Box plot shows the half width (hatched lines in trace 3) of the first and second VMN motoneuron action potentials (APs, for 8 neurons). Note the double spikelets that occur on first APs (inset in 4). b) Weak (black traces) and strong (blue traces) synchrony of vocal activity is reflected in vocal nerve compound potentials (top) and the voltage of intracellularly recorded VMN motoneuron APs (bottom). Electrical artefacts of midbrain stimulation are indicated by black arrowheads. Black arrows indicate spikelets. Inset shows magnification of unsynchronized motor output.c) Amplitude of VMN motoneuron afterhyperpolarization (AHP) and AP amplitudes are higher during fictive vocal behavior (red trace) than during antidromic stimulation (blue trace) via the vocal nerve. d) Current voltage responses of a VMN neuron show rapid AP adaptation. Inset shows color coded traces just before AP initiation (blue), with one (black) and two (red) APs. Note the decrease in second AP height in the red trace.

Across repetitions, the amplitude of the second VOC potential always exceeded the first (65.1 ± 15.3 μV vs. 36.6 ± 11.3 μV; n= 25 from five neurons; Mann-Whitney U test p<0.001) (Fig. 2a_3,4,5_). The reduced amplitude of the first VOC potential reflected a less synchronous and/or a partial activation of the motoneuron population. The first and last VOC potentials generally showed activity distributed over a broader time course (Fig. 2b; also see inset, amplifying end of vocal nerve record, for example, of “weak” synchrony). The amplitude of the first and last VOC potentials thus directly reflected the extent of synchronous motoneuron activation.

Intracellular motoneuron recordings showed broad depolarizations often present during the first and last VMN APs (Fig. 2a_3-5_, b). These depolarizations often displayed spikelets, strongly suggestive of asynchronous motoneuron activity (Fig. 2a_4_, b, c). A stronger afterhyperpolarization (AHP; −9.1 ± 2.2 mV below baseline [-61.8 ± 2.9 mV] levels; n=25 from five neurons) was detected with the first appearance of a VOC potential that was matched to an AP lacking the broad depolarization often observed with the first VOC potential.

Antidromic activation via the vocal nerve (electrodes implanted in vocal muscles; see Materials and Methods) revealed that motoneuron APs lacked the prominent AHP at their respective threshold (blue trace, Fig. 2c, right panel). This was consistent with intracellular square current injections showing no clear AHP after AP firing (Fig. 2d). Motoneuron AP and AHP amplitudes (relative to resting membrane potential) were significantly larger during VOC activity (AP: 41.50 ± 17.71 mV; n=39 for six neurons; AHP: −14.33 ± 7.84, n= 67 from six neurons) than following antidromic activation (AP: 28.09 ± 25.03 mV, AHP: −0.01 ± 1.04; n=59 from six neurons; Mann Whitney U test: AP p<0.001; AHP p<0.001; Fig. 2c).

The strong variability in amplitude of motoneuron APs and nerve potentials during a given VOC (e.g., Fig 2a_2-5_) was further reflected by differences in half-width of the first motoneuronal AP. Half-widths of the first APs riding on the aforementioned broad depolarizations were significantly wider than those of subsequent APs whose amplitude correlated with a higher amplitude in the corresponding VOC potential (first AP half width: 1.42 ± 1.07 ms; second: 0.72 ± 0.11 ms; n=40 from eight neurons; Mann-Whitney rank sum test: p<0.001). Larger VOC potential amplitudes thus indicated more extensive synchronous firing across the VMN population and correlated with narrower motoneuron APs.

### Motoneuron electrical coupling and after-hyperpolarization (AHP)

To investigate the origin of different motoneuronal AP and AHP amplitudes observed for antidromic-evoked APs and those during VOC activity, motoneurons were stimulated antidromically at varying amplitudes via the ipsilateral vocal nerve root (ad-ipsi, blue traces; Fig. 3a). At low amplitudes, we detected small depolarizations (Fig. 3a_2_) whose amplitude gradually increased with stimulation strength, i.e., with increasing recruitment of motoneuron axons and with shapes and peak latencies indicative of electrotonic coupling (3.7 ± 0.6 ms; n=30 from six neurons analyzed). Collision experiments using antidromic activated and intracellular evoked APs (via intracellular current injection) revealed these APs could not be blocked, i.e., they resulted from electrical coupling (Kiehn & Tresch, 2002; Pappas & Bennett, 1966) (Supp. Fig. 1).

**Figure 3:**
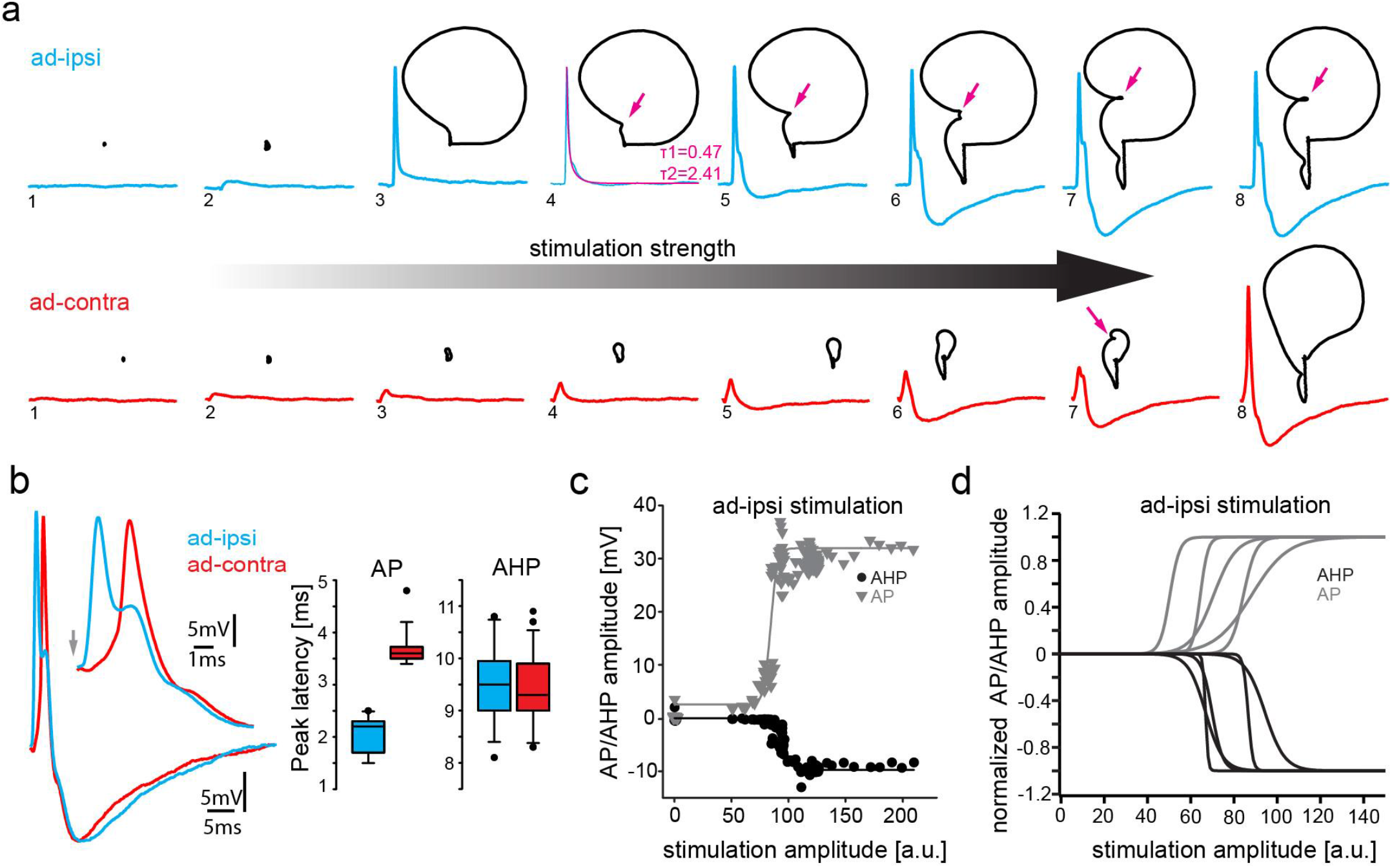
Antidromic stimulation activates motoneuronal afterhyperpolarization (AHP). a) Intracellular record of antidromically activated motoneuron upon ipsi- (blue) and contralateral (red) activation with increasing stimulation amplitude (schematized by big horizontal arrow). Black lines show phase plane plots and magenta arrows indicate additional depolarization prior to AHP onset. Magenta line in 4^th^ trial represents exponential fit for this recording, with the constants indicated. b) Overlay of ipsi- and contralateral stimulation of neuron shown in (a). Box plots show AHP and action potential (AP) peak latencies. c) AP (red) and AHP (blue) peak amplitude of VMN neuron response upon ipsilateral antidromic stimulation of variable amplitude (arbitrary units) and corresponding sigmoid fits (color coded lines). d) Normalized sigmoid fits of different neurons showing differences in recruitment threshold by antidromic stimulation, but similar time courses.

With increasing amplitude of vocal nerve stimulation, the axon of the respectively recorded motoneuron was eventually recruited and an antidromic AP invaded the recorded motoneuron as shown by the significantly shorter peak latency (2.0 ± 0.3 ms after stimulation; n=6) compared to the previously mentioned subthreshold depolarization (signed rank test: p<0.001) and the faster rise time (Fig. 3a_3_). These APs showed no AHP, but instead were characterized by a slow decay (double exponential fit; average time constant τ1: 0.85 ± 0.27; time constant τ2: 7.45 ± 3.17; n=21 from five neurons) back to the resting membrane potential (example shown in magenta trace in Fig 3a_4_). Surprisingly, an AHP started to appear with increasing antidromic stimulation amplitude (Fig. 3a_5_). The AP and AHP peak amplitudes increased with stimulation strength until each reached a plateau (Fig. 3a_6-8_, c, d). An AHP was never observed during intracellular square pulse current injections (Fig. 2d), raising the question on the origin of this AHP.

Phase plane plots of the membrane potential during ipsilateral antidromic activation revealed further changes in motoneuronal activity upon increasing stimulation amplitudes (Fig. 3a, black). The gradual appearance of the AHP, together with the absence of the AHP upon initial AP firing (Fig. 3a_3,4_), suggested that a further recruitment of motoneurons via the antidromic stimulation underlies AHP generation (Fig. 3A_5-8_). Phase plane plots further revealed an additional component: a broadening of the depolarization after AP firing (magenta arrows, Fig. 3A_4-8_). As this depolarizing component was not present at the recruitment threshold, it cannot have originated from the gap junction mediated coupling superimposed on the AP. In a few cases, this depolarizing component eventually led to a second AP firing (not shown).

A Renshaw cell-like recurrent inhibition, in which spinal motoneurons use an axon collateral to activate a local inhibitory circuit (Eccles et al., 1954; Renshaw, 1941), could be ruled out as the origin of the AHP given the lack of motoneuron axon collaterals (Fig. 1c) (Chagnaud & Bass, 2014) and the short onset of the AHP. To exclude that a motoneuronal axon collateral could arise at the periphery and enter via one of the nearby dorsal roots, the dorsal roots were bilaterally cut in two experiments. There was no difference in AHP amplitude (% of baseline) between cut and uncut recordings during VOCs (before cut: 17.04 ± 9.21, n= 150 from nine neurons; after: 15.19 ± 6.07, n=161 from nine neurons; Mann Whitney U-test: p=0.23; Supp. Fig. 2) or during antidromic stimulation (before cut: 12.37 ± 4.75, n=126 from nine neurons; after: 12.38 ± 4.47, n=127 from nine neurons; Mann Whitney U-test: p=0.59). These results excluded a motoneuronal collateral via one of the dorsal roots as the origin of the AHP.

Contralateral antidromic stimulation also revealed electrotonic potentials whose amplitude depended on stimulation strength (ad-contra, red traces; Fig. 3a). Electrotonically mediated potentials eventually reached threshold and evoked an AP. The peak latency of these APs was significantly longer (3.71 ± 0.36 ms; n=39 for six neurons) than ones elicited ipsilaterally (Fig. 3b; Signed rank test: p=0.001), while the peak latency of the subthreshold depolarization did not differ, consistent with their common origin from electrotonic coupling (see above). As tract tracing and intracellular neuron fills showed that motoneurons only innervate the ipsilateral muscle (Chagnaud & Bass, 2014), electrical coupling alone is thus able to drive AP firing, independent of whether motoneurons belong to the ipsilateral or contralateral VMN population.

As with the ipsilateral antidromic activation, an AHP component could clearly be distinguished in the contralateral antidromic stimulation experiments (Fig. 3a_4-8_). This AHP occurred independent of AP firing, emphasizing the independence of the two events within a given motoneuron.

Consistent with our findings in midshipman (Chagnaud et al., 2012), the ability to initiate an AP via electrotonic coupling was in strong contrast to our intracellular current injections that failed to initiate an AP in most cases, even at high current intensities (> 5 nA). In cases where intracellular current injection elicited an AP, motoneurons showed rapid adaptation of AP firing, likely due to weak somatic repolarization ability (see Fig. 2d). Gulf toadfish are, however, able to contract their vocal muscles for several hundred milliseconds (Fig. 1a). How can the muscle do this if motoneurons cannot fire for extended time periods due to the rapid AP adaptation seen during square pulse current injections? To test whether the AHP was required to de-inactivate motoneurons, we stimulated the motoneurons in which current injection led to AP firing with pulse trains of different frequencies. In contrast to long (> 50 ms) duration pulses (Fig. 2c), motoneurons showed no signs of AP adaptation to pulse trains with brief (< 5 ms) pulses, indicating the necessity of membrane repolarization for sustained motoneuron firing (Supp. Fig. 3). Stimulation was reliable into the behaviorally relevant physiological range (the pulse repetition rate/ fundamental frequency of toadfish vocalizations) with train frequencies tested up to 110 Hz. The weak repolarization capability and low excitability of the motoneurons thus provide the means to prevent sustained AP firing, which would decrease the extent of firing synchrony and precision across the VMN population.

### Network activity induces AHP

The presence of the AHP only at high antidromic stimulation amplitudes, i.e., high levels of motoneuron recruitment, strongly suggested a network-dependent activation of the AHP. To test this hypothesis, we ipsilaterally evoked an AP antidromically in a VMN motoneuron (ad-ipsi) at low threshold stimulation (i.e., without an AHP), followed by stimulation of the contralateral nerve (ad-contra), which resulted in a small electrotonic depolarization (Fig. 4a, b; also see supplemental movie 1). Subsequently, the delay of this second stimulation was reduced up to the time point of the first ipsilateral nerve stimulation (Fig. 4b). Once close to the antidromic AP, an AHP started to appear that increased in amplitude the closer the contralateral evoked potential came to the ipsilateral evoked antidromic AP (Fig. 4b; see heat map in 4a). These experiments suggested that an increase in overall depolarization in the vocal network is needed to generate the AHP. To test this hypothesis, we took advantage of the prominent, wide depolarization that often appeared in motoneurons at the end of a VOC (see Figs. 1d; 2a). Similar to the above, we moved an ipsi- or contralateral, antidromically evoked depolarizing potential into this depolarization occurring at the end of a VOC (ad-ipsi and ad-contra, Fig. 4c and d, respectively). With decreasing lag between the antidromically evoked potential and the depolarization during the VOC, an AHP started to appear in the contralateral evoked potential that increased in amplitude (Fig. 4c, d). This again showed the necessity of a network-wide depolarization in order to elicit the AHP.

**Figure 4:**
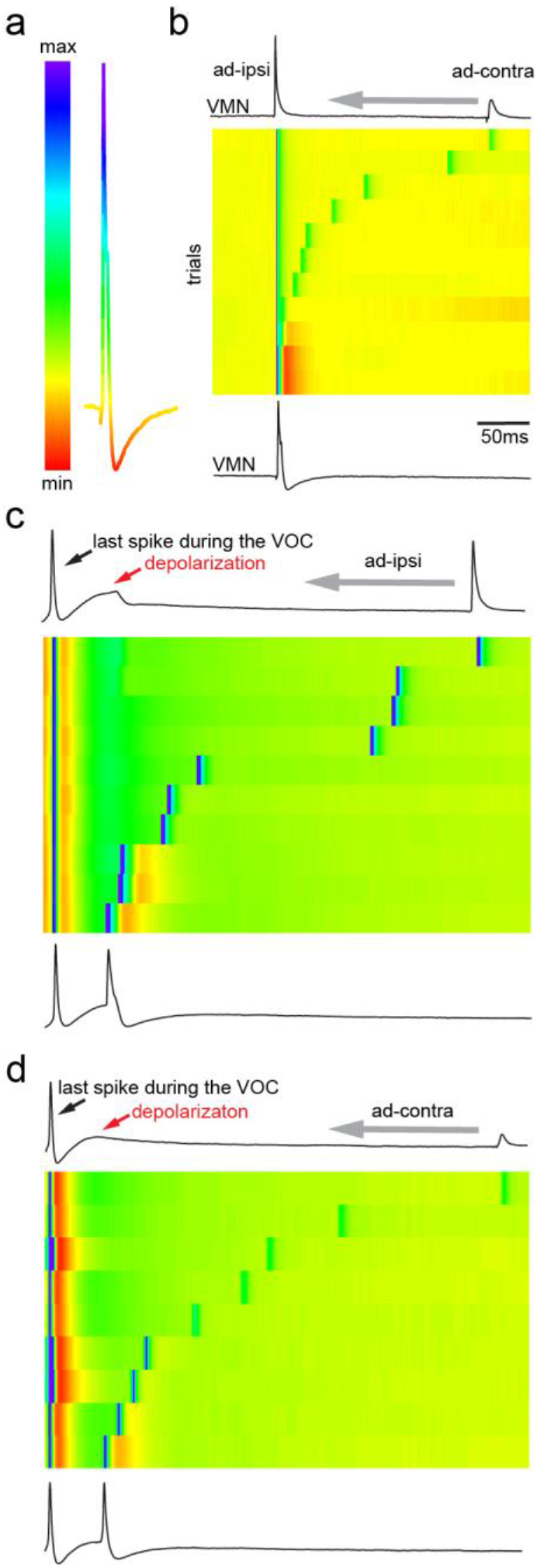
Antidromic activation of VMN motoneurons reveal network dependent afterhyperpolarization (AHP). a) antidromic evoked action potential color-coded to relative voltage amplitude and corresponding color bar. b) Stimulation dependent voltage matrix (SDVM; color code given in (a)) of ipsilateral antidromic activation followed by contralateral antidromic activation (that generated only a depolarizing potential) of decreasing latency reveals the appearance of an AHP when ipsilateral and contralateral stimulation overlap. In this situation ipsilateral antidromic stimulation was set to elicit an AP without an AHP. With decreasing distance of the contralateral stimulation an AHP started to appear (inset in SDVM). Top and bottom black lines represent first and last row of the SDVM in this and in following panels. c) SDVM showing last compound potential of a fictive vocalization (VOC) with associated depolarization (depol) and antidromic stimulation (set to elicit only a depolarization) of contralateral vocal nerve with decreasing latency. Contralateral potential generated an AP with decreasing distance to the depolarization that was accompanied by an AHP with further decrease in latency. d) As in F but with ipsilateral antidromic stimulation.

### Single motoneuron activity reflects population synchrony

To test the contribution of single motoneurons to network activity, we performed intracellular recordings of motoneurons using QX314 that blocks voltage-dependent sodium channels intracellularly (Yeh, 1978). After intracellular iontophoresis of QX314, motoneurons exhibited about a 30% decrease in AP amplitude compared to baseline conditions during VOCs (baseline: 31.3 ± 5.1 mV; QX314: 25.5 ± 5.9 mV; Sign ranked test: p=0.002) (Fig. 5a). There was a much more prominent, about 70%, decrease in antidromically evoked AP amplitude relative to baseline (baseline: 19.1 ± 11.9 mV; QX314: 5.9 ± 4.0 mV;n=35 from 7 neurons; Sign ranked test: p<0.001) (Fig. 5b). While a significant difference in the AHP during VOCs was observed (baseline: −9.4 ± 2.0 mV; QX314: −3.2 ± 2.0mV; n=35 from 7 neurons; paired t-test: p<0.001), no significant change could be detected in the AHP following antidromic activation (baseline: −2.8 ± 4.1mV; QX314: −2.9 ± 4.0mV; n=35 from 7 neurons; signed ranked test: p=0.62) (Fig. 5a, b). These experiments demonstrated that (i) the contribution of the recorded motoneuronal AP firing to the firing of that motoneuron is rather small during VOC activity (the activity of the neuron is dominated by gap junction coupled potentials), (ii) during a VOC, the AHP amplitude of the recorded neuron only partly depends on its firing an AP (also see above), and (iii) most of the activity displayed by a given motoneuron reflects population-level motoneuronal activity.

**Figure 5:**
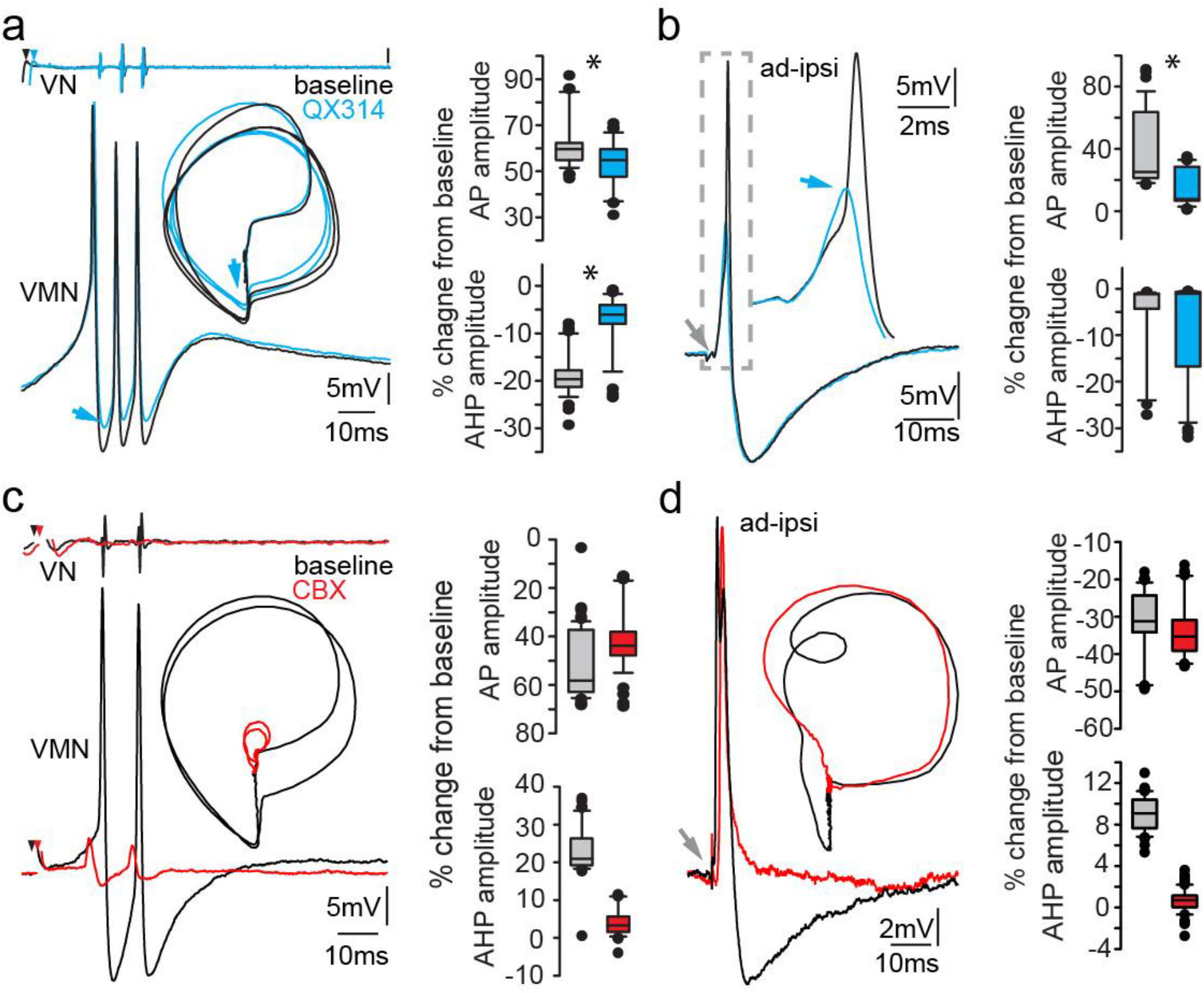
Network activity is essential to generate the motoneuron afterhyperpolarization (AHP). a, b) Intracellular recording of VMN neuron action potentials (APs) with QX314-filled electrodes during fictive vocal activity (a) and ipsilateral antidromic stimulation (b). Inset in A shows phase plane plot and in (b) higher magnification of antidromically evoked APs. Box plots show %change from baseline level for AP and AHP during either condition. Blue arrows point to the differences in AP and AHP levels. c, d) Intracellular recording of one VMN neuron before (black) and one VMN neuron after (red) blocking gap junctions by application of carbenoxelone (CBX). Inset shows phase plane plot. Box plots show % change from baseline level for AP and AHP during either condition.

### Necessity of gap junctional coupling for AHP activation

Having observed that the AHP is highly dependent on activation of the VMN population and not on single neuron AP firing (Figs. 3–5a,b), we next blocked gap junctional coupling to determine if the AHP originated from a network activation. A combined superfusion of the exposed vocal hindbrain region coupled with pressure injection of carbenoxolone (CBX, a gap junction blocker) directly into VMN severely impared the vocal network’s ability to generate synchronized motor discharges as evidenced by barely detectable VOC activity (Fig. 5c, red trace). However, low amplitude VOC-related activity could still be detected in intracellular recordings from motoneurons upon midbrain stimulation, showing that the vocal network could still be activated (Fig. 5c). Upon antidromic stimulation (Fig. 5d), APs had a similar amplitude as during control conditions, showing that the loss of vocal-related activity during midbrain activation was not due to CBX impairment of motoneuron action potential-generating capacity, but to decreased electrotonic input to the motoneurons (% change from baseline: control: −31.96 ± 9.31; CBX: −33.67 ± 9.68; n=54 from five neurons; Mann Whitney U test: p=0.079). No AHP could be elicited in antidromic activated VMN neurons. This showed that gap junctional coupling is indeed required to activate the AHP.

### Dependence of AHP activation on inhibitory input to VMN

Having identified that gap junctional coupling was essential for the AHP, we next tested whether inhibitory input could provide a source of this AHP.

#### Bicuculline injections into VMN

Injections of bicuculline, a potent GABA_(A)_ receptor antagonist, into VMN led to an increased rise time of the onset depolarization and in AP amplitude during VOCs (Fig. 6 a,b; baseline % change; control: −34.38 ± 15.42, n=121 from ten neurons; bicuculline: −45.73 ± 14.58, n=177 from thirteen neurons; Mann Whitney U test: p<0.001) and a decrease in AHP amplitude (baseline % change; control: 18.23 ± 26.32, n=121 from ten neurons; bicuculline: −6.56 ± 8.88, n= 177 from thirteen neurons; Mann Whitney U test: p<0.001). However, bicuculline did not abolish the AHP. Motoneurons were also still able to fire APs during VOCs, although the amplitude of VOC potentials was on average reduced by more than half (control: 92.77 ± 16.70 μV, n=96 from six neurons; bicuculline: 44.85 ± 8.39 μV, n=155 from six neurons; Mann Whitney U test: p<0.001), indicating an effect of inhibition on the extent of synchrony throughout the VMN population.

**Figure 6:**
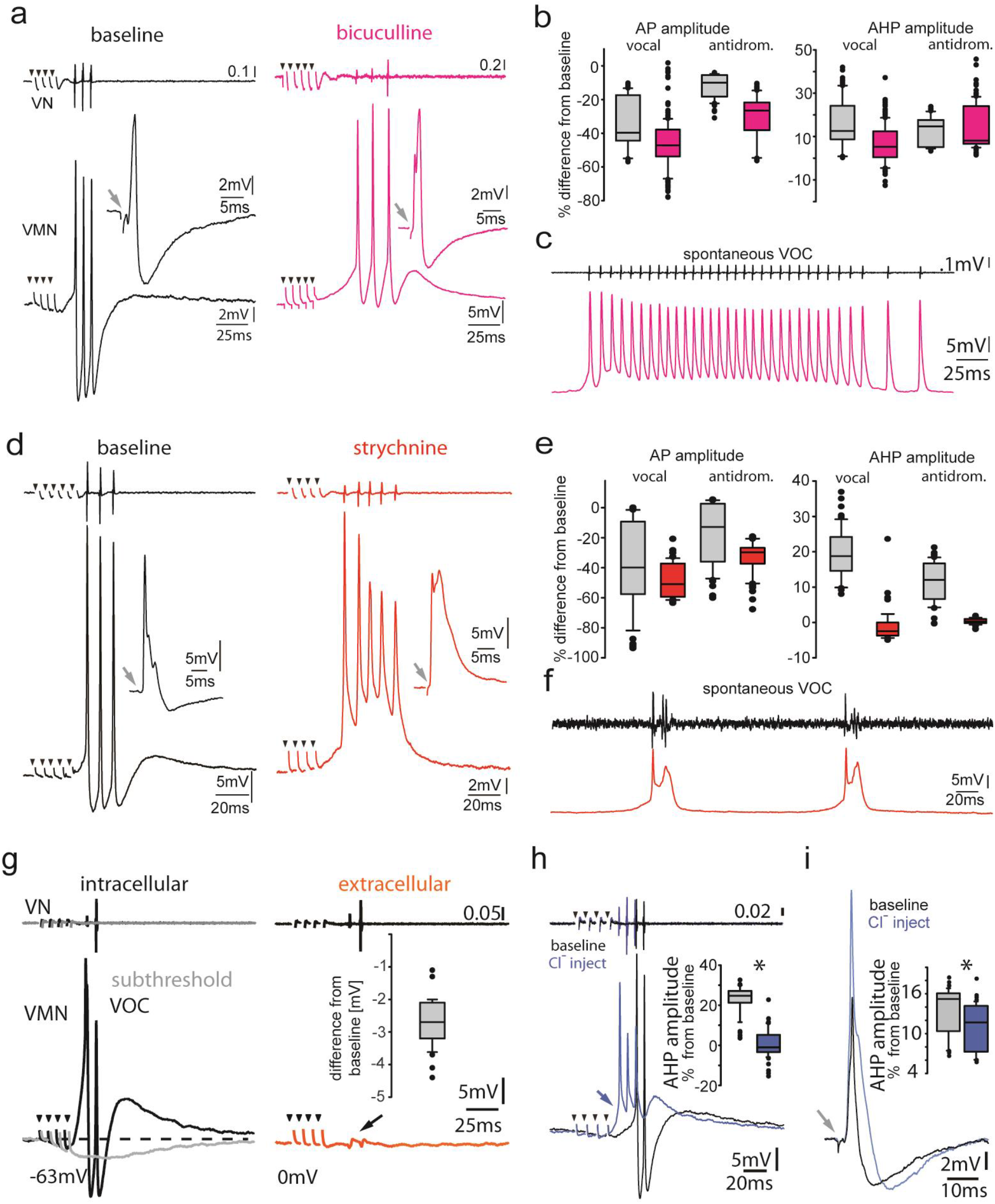
Inhibitory input is essential for vocal patterning. a) Intracellular recordings from a motoneuron before (left) and from another after (right) bicuculline injection into VMN. Insets show antidromic evoked action potentials for the respective neurons. b) Box plots showing the change in amplitude of the action potential (AP) and of the afterhyperpolarization (AHP) for control (gray) and bicuculine injected(magenta). c) Intracellular recordings of a motoneuron (magenta trace) during a spontaneous fictive vocalization (VOC, black trace). d) Intracellular recordings from a motoneuron before (black) and another motoneuron after (red) strychnine injection into VMN. Insets show antidromic evoked action potentials for the respective neurons. e) Box plots showing the change in amplitude of the action potential (AP) and of the afterhyperpolarization (AHP) for control (gray) and strychine injected (red). f) Intracellular recordings of a motoneuron (red trace) during two spontaneous fictive vocalization (VOC, black trace). g) Intracellular recording of VMN motoneuron during midbrain subthreshold stimulation shows decrease in membrane potential (gray trace) compared to during vocal activity (black trace) and extracellular-recorded activity (red trace). h,i) Intracellular chloride injection into one motoneuron to modify chloride reversal potential shows prominent difference in amplitude during vocal activity (h), but not during antidromic stimulation (i). Box plots show change in amplitude of afterhyperpolarization (AHP; percent change form baseline levels).

The amplitude of the antidromically evoked motoneuron AP increased (baseline % change; control: −11.37 ± 7.08, n=74 from 5 neurons; bicuculline: −29.97 ± 12.61, n= 98 from seven neurons; Mann Whitney U test: p<0.001), but no significant difference was found in the antidromically evoked AHP (baseline % change; control: 12.50 ± 6.63, n=74 from 5 neurons; bicuculline: 13.97 ± 10.13, n= 98 from seven neurons; Mann Whitney U test: p=0.306; Fig. 6a inserts, 6b). With time, spontaneous VOCs appeared (Fig. 6c) that showed similar decreases in motoneuron AHP amplitude not seen in spontaneous VOCs without prior bicuculline injections). These results are consistent with dense GABAergic input to the Gulf toadfish VMN (Rosner et al., 2018). While this input affects the motoneuron AHP, it only partially contributes to its generation.

#### Strychnine injections into VMN

To test the influence of glycinergic input on generation of the AHP, we pressure injected the glycine receptor antagonist strychnine into the VMN. Immediately after strychnine injection, there was an increase in the rise time of the onset depolarization in motoneurons during midbrain-evoked VOCs (Fig. 6d; resembles bicuculline effect, 6a). The AHP of motoneurons also disappeared (baseline % change; control: 19.31 ± 7.14, n=62 from five neurons; strychnine: −1.15 ±4.65, n=48 from five neurons; Mann Whitney U test: p<0.001; Fig. 6d, e). Similarly, and in contrast to the bicuculline experiments, the AHP during antidromic stimulation was completely abolished (baseline % change; control: 11.14 ± 5.89, =62 from five neurons; strychnine: 0.34 ± 0.79, n=48 from five neurons; Mann Whitney U test: p<0.001; Fig. 6d inserts, e) and accompanied by a longer depolarization of the antidromic AP (Fig. 6d inserts). These results suggested that glycinergic neurons, activated via gap junctional coupling, were responsible for generating the AHP during antidromic stimulation.

Following strychnine injections, spontaneous VOCs appeared with similar decreases in motoneuron AHP amplitude not seen for spontaneous VOCs under control conditions without prior strychnine injections (Fig. 6f). Since VMN motoneurons lack axon collaterals (Fig. 1c), the results imply that gap junction coupling between motor and glycinergic VPN neurons is sufficient to drive AP firing of glycinergic neurons.

Typically, VOCs show sharp peaks and stable though sometimes variable intervals between successive potentials, similar to the sound pulses within natural grunts (e.g., (Maruska & Mensinger, 2009; McIver et al., 2014). This was also the case for spontaneous VOCs following GABA injections (Fig. 6c). Visual inspection of spontaneous VOCs following strychnine injections suggested a loss of this stereotypy; double peaked potentials varying in width were reminiscent of VOCs associated with weak VMN synchronization (Fig. 2b). Given strychnine and GABA were injected throughout the VMN, we measured interpulse intervals for midbrain-evoked VOCs that reflect VMN population activity. This analysis revealed a significant difference for the first and second VOC intervals between control conditions and following strychnine application (control: 8.21 ± 1.30 ms, n=73 from 5 neurons; strychnine: 7.80± 2.46 ms, n=56 from 4 neurons; Mann Whitney U test: p=0.037; spontaneous VOCs were too few in number to warrant measurement). Thus, strychnine decreased interspike intervals across the VMN population (also compare 6a and 6d) and hence VOC frequency. Variability also increased significantly, with strychnine injected VOC intervals being more variable (Levene’s test: F=13,866, p<0.01).

#### Intracellular chloride injections into vocal motoneurons

Having identified that blocking inhibition was essential for the AHP, we next tested whether the effect of inhibition was detectable in individual neurons. At sub-threshold midbrain stimulation levels to induce a VOC compound potential, tonic membrane hyperpolarizations (on average, −2.69 ± 0.73 mV below resting membrane potential; n=44 from five neurons) could be detected (Fig. 6a, gray trace, subthreshold, blue trace suprathreshold), indicating the activation of inhibitory inputs. To test whether these hyperpolarizations were artefacts originating from electrical stimulation, we retracted the electrode from the intracellular space (Fig. 6g, orange trace). This abolished the hyperpolarizing components. Due to the high synchrony of motoneuron APs during vocal activity, field potentials could be detected in the nerve even with our high resistance electrodes (Fig. 6g, black arrow). Changing the chloride reversal potential in the respective neurons by intracellular chloride injections via 3M KCl-filled electrodes revealed a prominent inhibitory input to motoneurons during VOCs as the membrane potential showed significant changes in the degree of repolarization compared to baseline levels (Fig. 6h, blue arrow). The AHP during VOCs was heavily reduced (Fig. 6h, blue trace; % change from baseline; baseline: 21.2 ± 8.7; chloride injected: 2.6 ± 11.9; n=51 from six neurons; paired t-test: p<0.001), thus indicating an inhibitory contribution to vocal behavior. Antidromic stimulation still showed the AHP, however at a reduced amplitude (Fig. 6i; baseline: 13.13 ± 3.72 mV; chloride injected: 10.64 ± 3.80 mV; n=45 from four neurons; paired t test: p<0.001). It should be noted that despite the short time difference between the VOC and the antidromic stimulation (<400 ms), a much smaller effect was detected on the antidromic AHP. This seemingly contradictory result is likely due to the electrotonic coupling of the network where other motoneurons contribute to the activity of individual motoneurons. Manipulation of only one out of the hundreds of motoneurons in VMN via chloride injections is thus not sufficient to reveal the full extent of inhibitory activity.

#### Vocal premotor neurons are excited by gap junctional coupling

Antidromic stimulation of the vocal nerve leads to membrane depolarizations (via electrotonic input) in premotor VPN neurons in the closely related midshipman (Bass & Baker, 1990). This, together with the greater excitability of VPN neurons (Chagnaud et al., 2011), could potentially generate a feedback activation of glycinergic neurons. We recorded from toadfish VPN neurons to test whether gap junctional coupling is in fact able to induce AP firing in the VPN population, which would be needed to recurrently activate or inhibit motoneurons. Toadfish VPN neurons generate two depolarizing components, one before and one during VOCs that might reflect motoneuron activity leaking through gap junctions (Fig. 7a) (Chagnaud & Bass, 2014). During antidromic stimulation of the vocal nerve, VPN neurons showed small subthreshold potentials similar to VMN neurons indicative of gap junctional coupling, as well as APs at higher stimulation amplitudes (Fig. 7b). Thus, antidromic activation in motoneurons could elicit AP firing in VPN premotoneurons. The latency of these APs (5.05 ± 0.48 ms, n=144 from 6 neurons) roughly coincided with the depolarization following antidromically elicited APs in motoneurons (Fig. 3a, magenta arrow).

**Figure 7:**
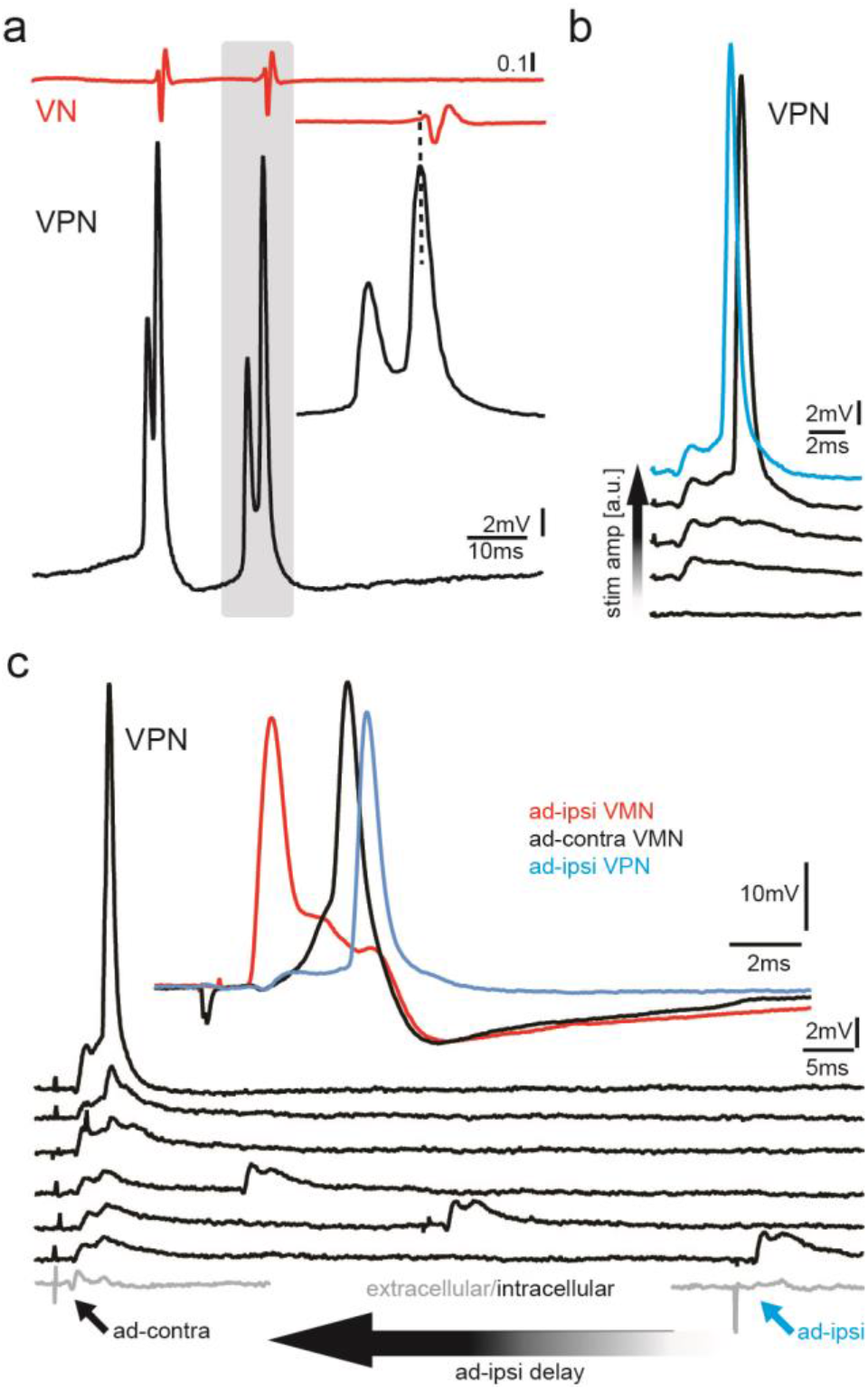
Premotoneurons can be activated via gap junctional coupling. a) Intracellular recording of VPN neuron during fictive vocal activity show the characteristic double spiking of VPN neurons. Inset shows timing between vocal nerve (VN) and VPN neuron activity. b) Waterfall plot of intracellular VPN recordings during ipsilateral antidromic vocal nerve stimulation leads to depolarizing potentials in VPN neurons that eventually lead to AP firing with increasing stimulation amplitude. c) Intracellular recording of one VPN neuron during ipsilateral (blue) or contralateral (red) antidromic vocal nerve stimulation. d) Waterfall plot of intracellular recordings from a VPN neuron during antidromic stimulation of the ipsi (ad-ipsi) and contralateral (ad-contra) vocal nerve. With decreasing latency between the ad-contra and the ad-ipsi stimulation (at identical stimulation amplitudes) an action potential can be elcitied, showing the capability of gap junctional coupling to induce AP firing. Inset shows recordings from a VMN neuron during antidromic stimulation of the contralateral and ipsilateral side and from a VPN neuron during ipsilateral antidromic stimulation (not simultaneously recorded).

To test if a network component might also be important to activate VPN neurons, we stimulated the ipsilateral and contralateral vocal nerve (at varying latencies) to antidromically generate subthreshold membrane depolarizations in VPN neurons. A decrease in latency between the ipsilateral and contralateral stimulation pulses eventually led to AP firing in VPN neurons, thus showing that AP firing in VPN neurons also depends on gap junctional activation of the vocal network (Fig. 7c). VPN activation likely led to motoneuron depolarization (Fig. 3, magenta arrows) and activation of gap junction coupled glycinergic neurons (Rosner et al., 2018) that, in turn, inhibit motoneurons and generated the AHP.

#### Glycinergic boutons contact vocal motoneurons

We previously identified a subpopulation of glycinergic VPN neurons, suggesting direct glycinergic input onto motoneurons (Rosner et al., 2018). To confirm and identify the location of glycine release onto vocal motoneurons, we used multi-color super-resolution structured illumination microscopy (SR-SIM) imaging, which can achieve an ehanced resolution of ~100 nm in the lateral (x-y) and ~300 nm in the axial (z) planes (Gustafsson, 2008). SR-SIM imaging demonstrated dense, punctate glycinergic-immunoreactive labeling throughout VMN. Overlap of glycinergic signal with labelling for synaptic vesicle protein (SV2) revealed prominent boutons directly abutting motoneuron somata (Fig. 8a-c). Glycinergic boutons were also observed on motoneuron dendrites within the contralateral VMN (Fig. 8d-h) and VPN (Fig. 8d, i-k). These results demonstrate an anatomical basis for glycinergic release and inhibition spatially distributed across the entire somato-dendritic extent of vocal motoneurons.

**Figure 8:**
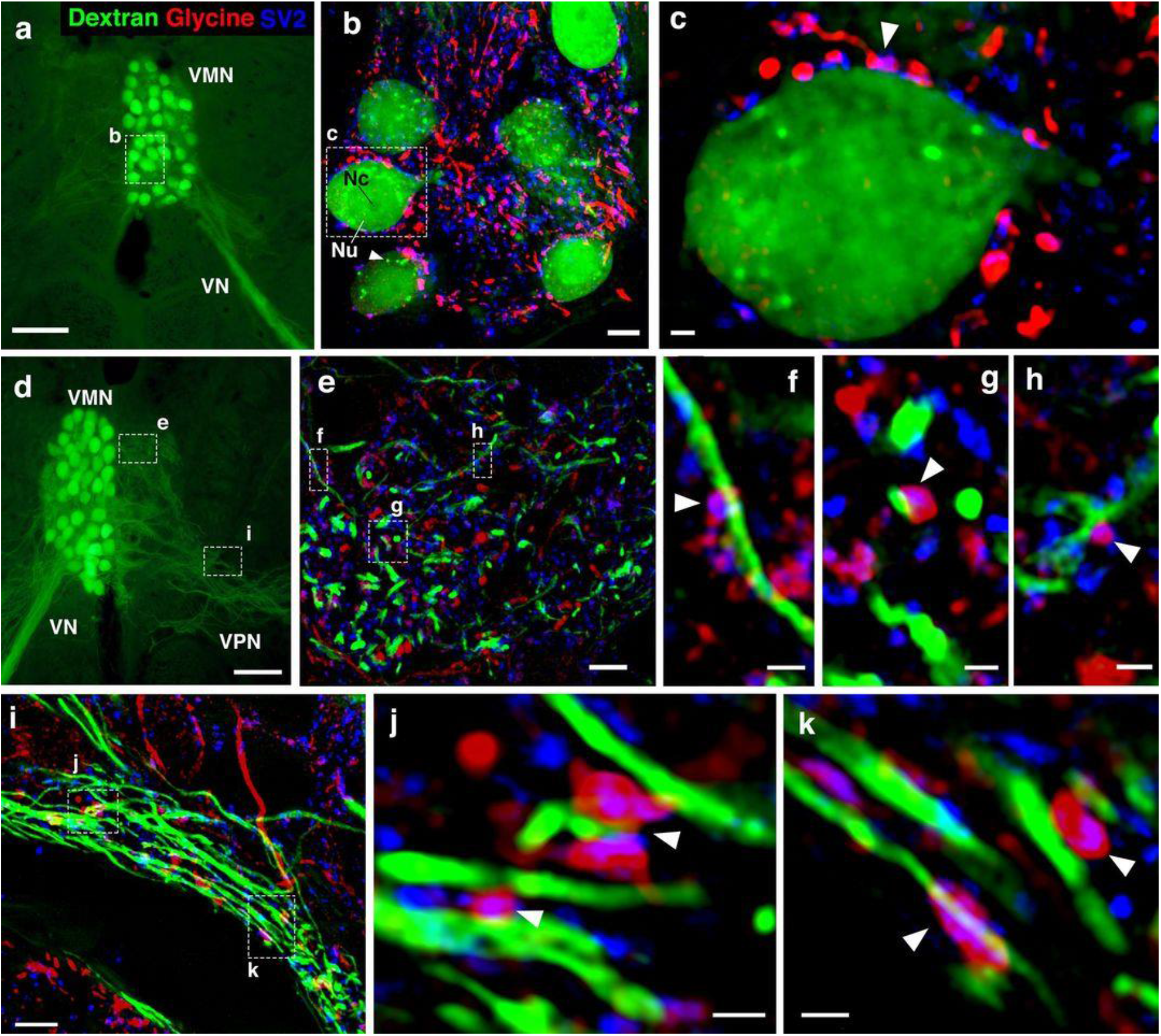
Glycinergic input on motoneuron somata and dendrites. a, d) Widefield micrographs show dextran-filled motoneurons (green) in vocal motor nucleus (VMN) of two different fish. Dextran labeling of one vocal nerve (VN) results in filling of ipsilateral VMN somata and dendrites that extend into contralateral VMN (d, box e) and bilateral into adjacent VPN columns (d, box i; also see Fig. 1c). White boxes correspond to maximum projections of super-resolution structured illumination microscopy (SR-SIM) z-stacks (b, e, i) that show glycine (red) and synaptic vesicle protein 2 (SV2, blue) immunoreactive label in proximity to VMN somata (b) and dendrites (e, i). b) VMN somata are surrounded by dense glycinergic and SV2 puncta. Large nucleus (Nu) with nucleolus (Nc) is evident in several somata; punctate dextran labelling within cytoplasm (white arrowhead) likely indicates sequestration of dextran into vesicles. c) A single optical section (0.1 μm) from SR-SIM z-stack in (b) shows a glycinergic fiber forming a bouton (white arowhead) on a motoneuron soma. Overlap of glycine and SV2 (magenta, white arrowhead) indicates a site of neurotransmitter release, and a likely synapse. Additional optical sections show glycinergic release sites (white arrowheads) on motoneuron dendrites extending into contralateral VMN (f, g, h, corresponding to boxes in e) and bilaterally into VPN columns (j, k, corresponding to boxes in i). Scale bars represent 100 μm in widefield micrographs (a, d), 5 μm in SR-SIM maximum projections (b, e, i) and 1 μm in SR-SIM optical sections (c, f, g, h, j, k).

## Discussion

Our experiments show that gap junction-mediated activation of glycinergic neurons, leading to feed-forward inhibition, can account for extreme temporal precision in motoneuron firing and likely also contributes via gap junctional feedback to pattern generation. The level of temporal fidelity in this network is perhaps rivalled only by that of the electromotor system of teleosts. Although electronic coupling and intrinsic neuronal properties seem to play a predominant role in temporal precision in that system, other mechanisms have not been fully explored (Bennett, 1971; Moortgat et al., 2000a, 2000b; Moortgat et al., 1998). The concurrent activation of neurons required for rapid and precise activation of muscle groups underlying acoustic signaling in fishes (Chagnaud et al., 2012) and tetrapods (Kwong-Brown et al., 2019; Mead et al., 2017), might all benefit from feed-forward inhibition. The extent of temporal precision across an entire neuronal population generated in toadfish far exceeds the magnitude of synchronization known for other vocal circuits where motoneurons are gap junctionally coupled to premotoneurons (Barkan & Zornik, 2019). More broadly, this system may provide further insight into the fine temporal control of motor patterning and activation of neuronal populations found throughout the central nervous system (Kros et al., 2017; Llinás, 2014; Song et al., 2016).

### Electrical coupling and inhibition in a vocal network

Electrotonic coupling is known to increase the synchronous activity of neuronal populations (Alcamí & Pereda, 2019). An important role of gap junctional coupling in the toadfish vocal network is its involvement in generating the prominent AHP in motoneurons detected during both midbrain-evoked VOCs and antidromic motoneuron stimulation. This hyperpolarization, which could not be elicited upon intracellular current injection, is essential to generating a highly synchronous motor output that underlies the ability to generate vocalizations with extreme temporal precision.

Toadfish vocal motoneurons receive both GABAergic and glycinergic inputs (Rosner et al., 2018). The present report strongly suggests a tonic inhibitory input during vocal behavior mediated by GABA and a phasic glycinergic input phase-coupled to the VMN-VPN firing pattern due to electrotonic coupling of glycinergic neurons to motoneurons. We previously described GABAergic inhibition at the single cell and network levels in midshipman (Chagnaud et al., 2012). The glycinergic component, confirmed here with SR-SIM, provides a feed-forward inhibition from premotoneurons to motoneurons. This is consistent with the timing of inhibition during vocal activity, in which the latency of the inhibitory potential is shorter than during antidromic activation. Blocking GABAergic action in VMN resulted in greatly reduced VOC amplitude, reflecting loss of VMN synchrony. Blockage of glycine action alone abolished the AHP that was coupled to increased VOC frequency and reduced stereotypy. A distinguishing temporal feature between grunts and boatwhistles is the greater stability of time intervals between successive sound pulses in boatwhistles (Maruska & Mensinger, 2009), which is also the case for the agonistic grunts and boatwhistle-like advertisement “hum” of midshipman (Brantley & Bass, 1994; McIver et al., 2014). The results strongly imply a salient role for glycinergic premotor neurons in determining this temporal feature of toadfish calls that varies with social context (aggression versus courtship). Interestingly, for fish of both sexes collected during the non-breeding season, the AHP did not appear during antidromic stimulation, although it was present during the VOC (not shown). While we did not quantify this result, a seasonal variability in the expression of the AHP upon antidromic stimulation appeared to be present. This variation might be due to the previously observed seasonal variability in expression levels of connexin transcripts in the closely related midshipman fish (Feng et al., 2015).

Antidromic activation is a highly artificial condition in which depolarizations are first elicited in VMN, and then transmitted via gap junctions to VPN. Thus, a recurrent inhibitory pathway is activated during antidromic stimulation. During natural vocal behavior, VPN would be feeding forward to VMN, given that VPN fires before VMN during VOCs (Bass & Baker, 1990; Chagnaud et al., 2011; Chagnaud & Bass, 2014). Consequently, excitatory VPN neurons likely activate VPN’s glycinergic neurons before the motoneurons. Electrotonic coupling of VPN excitatory neurons to glycinergic VPN neurons thus activates the latter, which likely causes feed-forward inhibitory action upon motoneurons. Electrophysiological recordings from VPN glycinergic neurons during vocal activity are needed to verify this hypothesis. If proven, we expect these neurons to fire rhythmically like other VPN neurons directly coupled to VMN (Chagnaud et al., 2011; also see Fig 7a). As gap junctions appear to be birectional, we cannot entirely rule out the recurrent activation hypothesis (Alcamí & Pereda, 2019).

Our antidromic stimulation experiment suggests a feedback loop of glycinergic inhibitory neurons activated upon by depolarization of the VMN-VPN network. However, the shorter latency of the inhibitory potential during vocalization compared to antidromic activation argues against a recurrent inhibition hypothesis and favors a feed-forward hypothesis. In any case, inhibitory action can serve multiple functions. First, it repolarizes motoneurons, a feature essential to VMN’s ability to repetitively fire APs, as seen by our pulse train experiments. Second, it generates a window of decreased excitability that prevents motoneuron misfiring outside of the VPN rhythm (due to the long time course of inhibitory action compared to the normal activation rhythm of ca. 100 Hz; see supplemental movie 2). This is essential to the ability of the next incoming excitatory input of rhythmically firing VPN neurons to set the firing frequency of vocal motoneurons. Hence, this gap junction mediated, feed-forward glycinergic inhibition can account for the ultra-precise temporal patterning in the millisecond time range that characterizes the vocal network and behavior of toadfishes (Bass & Baker, 1991; Chagnaud et al., 2011; Chagnaud et al., 2012; Kéver et al.; Pappas & Bennett, 1966). A similar mechanism has previously been shown for the Mauthner cell escape circuit(Faber et al., 1989; Furukawa & Furshpan, 1963; Korn & Faber, 2005; Sillar, 2009; Zottoli & Faber, 2000), and may operate in other vocal systems in teleosts and tetrapods that are dependent on comparable levels of temporal precision (Bass et al., 2015; Kwong-Brown et al., 2019; Mead et al., 2017; Nelson et al., 2018; Rome, 2006).

### Consequence of motor-premotor coupling

The notion that vertebrate motoneurons are only passive components has recently changed. Motor-premotor coupling in other systems (Barkan & Zornik, 2019; Falgairolle et al., 2017; Lawton et al., 2017; Matsunaga et al., 2017; Song et al., 2016) are in line with our observation that motoneuron activity has substantial impact on premotor patterning. While we do not directly show the effect of motoneuronal activity back-propagating through gap junctions on the VPN firing pattern, we hypothesize that the characteristic “pacemaker” potential that leads to VPN oscillatory-like activity in midshipman (Chagnaud et al., 2011) originates from motor-premotor coupling. This would mean excitatory potentials from the VMN network are shunted by the presumed feed-forward inhibitory action, while the inhibitory potentials would be allowed to travel through gap junctions to contribute to VPN patterning (e.g., see Fig. 7a; also see Chagnaud et al., 2011). Soma-dendritic recordings from VPN are needed to test the contribution of motoneurons to their pattern generating ability. It is, however, conceivable that the gap junctional coupling between VPN and VMN neurons also affects VPN activity and is another example of how premotor and motor neurons interact to generate motor patterns (Barkan & Zornik, 2019).

### Why extreme temporal precision?

From a behavioural perspective, there are several reasons why there might be strong selection for vocal behaviors with extreme temporal precision that is directly determined by a central timing circuit, in this case the VPN-VMN network. One reason is that call types differing in temporal properties specify behavioral state. For example, broadband grunts indicate an aggressive state during defense of resources such as nest sites, whereas mutliharmonic advertisement calls are courtship signals indicating readiness to mate (for reviews, see Bass & McKibben, 2003; Ladich et al., 2006). Underwater playbacks show that both male and female toadfishes, including Gulf toadfish, distinguish call types and interpulse intervals (Fish, 1972; McKibben & Bass, 1998; McKibben & Bass, 2001; Remage-Healey & Bass, 2005; Winn, 1972). Individual differences in boatwhistle properties are linked to male reproductive success in toadfish (Vasconcelos et al., 2012). More broadly, behavioral evidence supports a role for interpulse intervals in individual and species recognition in other soniferous teleosts (Amorim et al., 2015; Gerald, 1971; Maruska et al., 2007; Myrberg & Riggio, 1985; Myrberg & Spires, 1972). From a sensory-motor perspective, the auditory system of fishes is exquisitely adapted to encoding the temporal properties of individual calls, including PRR and fundamental frequency (Bass & McKibben, 2003; Fay & Edds-Walton, 2008). As Capranica wrote (Capranica, 1992), the vocal and auditory systems “co-evolved and we should expect them to share the same underlying code for signal generation and recognition”.

From an environmental perspective, transmission distance is limited by the frequency content of calls in shallow water habitats like those where toadfishes defend nests and engage in acoustic courtship (Bass & Clark, 2003; Fine & Lenhardt, 1983; Gerald, 1971). The principal limitation to call PRR and fundamental frequency in species like toadfishes that generate calls with frequency content mainly ≤500 Hz is the contraction rate of muscles that, in this case, drive swim bladder vibration. To overcome such limitations and enhance transmission distance, toadfishes adopted superfast muscles (Elemans et al., 2014; Nelson et al., 2018; Rome, 2006) driven by a central network that directly determines extreme temporal precision in the synchronous activation of muscle fibers to maximize amplitude as well as PRR and fundamental frequency. While a combination of mechanisms increase temporal precision in the time domain, gap junction-mediated, feed-forward glycinergic inhibition provides a novel means to achieve extreme temporal precision in motor coding at a network level.

## Materials and methods

### Animals

Gulf toadfish of both sexes (n = 18; standard length: 6 females: 11.7-13.6 cm; 12 males 15.3-25.7 cm) were obtained from a commercial source (Gulf Specimen, Panacea, Fla., USA) and housed in saltwater aquaria in an environmental control room held at 22°C on a 14:10 light: dark cycle. Though field studies mainly characterize male vocalization, both sexes are capable of acoustic signaling (e.g., Demski & Gerald, 1972; Fine & Thorson, 2008; Remage-Healey & Bass, 2005). All experimental methods were approved by the Cornell University Institutional Animal Care and Use Committee.

### Surgery for neurophysiology

Surgical and recording methods (see below) were adopted from prior studies (Bass & Baker, 1990; Chagnaud et al., 2012). Animals were deeply anesthetized during all surgical procedures [immersion in aquarium water containing 0.025% benzocaine (ethyl p-amino benzoate); Sigma, St. Louis, Mo., USA]. For antidromic stimulation of VMN motoneurons, bipolar silver wire electrodes insulated with enamel except at the tips (0.15 mm diameter; 0.3 mm between tips) were implanted at the level of the swim bladder between each muscle and the bladder wall, immediately adjacent to the vocal nerve. A dorsal craniotomy was performed to expose the brainstem and the paired occipital nerves that give rise to the vocal nerve (Fig. 1b). After surgery, animals received an intramuscular injection of bupivacaine anesthetic (0.25%; Abbott laboratories, Chicago, Ill., USA) with 0.01 mg/ml epinephrine (International Medication Systems, El Monte, Calif., USA) near the wound site and then an intramuscular trunk injection of the muscle relaxant pancuronium bromide (0.1–1 μg/g of body weight); bupivacaine was administered every four hours until euthanasia. Animals were placed in a plexiglass tank and perfused over the gills with artificial seawater at 18–20 °C, the same temperature as their home aquarium water.

### Monitoring and activation of vocal motor behavior

Teflon-coated, silver wire electrodes (75 μm diameter) with exposed ball tips (50–100 μm diameter) were used to record the vocal motor volley, hereafter referred to as a fictive vocalization (VOC), from the vocal nerves that innervate the vocal muscles attached to the swim bladder. Signals were amplified 1,000-fold and band-pass filtered (300–5 kHz) with a differential AC amplifier (Model 1700, A-M Systems). VOCs were evoked by current pulses delivered to vocal midbrain areas via insulated tungsten electrodes (5 MΩ impedance; A-M Systems). For display purposes, electrical artefacts were truncated in the illustrations (marked by black arrowheads in Figs. 1, 2, 5, 6). Current pulses were delivered via a constant current source (model 305-B, World Precision Instruments). A stimulus generator (A310 Accupulser, World Precision Instruments) was used to generate TTL pulses with a standard stimulus of 5 pulses at 200 Hz. Each pulse train equaled one stimulus delivery with inter-stimulus intervals of 1 s. During recordings, inter-pulse intervals (100–300 Hz) and total pulse number (2–10) varied. Occasionally, VOCs also occurred spontaneously.

### Neurophysiological recordings

Glass micropipettes (A-M systems) for intracellular recordings were pulled on a horizontal puller (P97, Sutter Instruments) and were filled with either a 5% neurobiotin solution in 0.5 M KCOOH (resistance 35–50 MΩ) or with 2 M KCOOH. Neuronal signals were amplified 100-fold (Biomedical Engineering) and digitized at a rate of 20 kHz (Digidata 1322A, Axon Instruments/Molecular Devices) using pCLAMP 9 software (Axon Instruments). An external clock (Biomedical Engineering) sending TTL pulses was used to synchronize stimulus delivery and data acquisition. Electrode resistance was monitored while searching for neurons by a current step applied to the recording electrode. In some cases, small amounts of negative current were applied to stabilize the membrane potential after penetration.

### Pharmacological manipulations

The dorsal roots at the level of the vocal occipital roots were cut with iridectomy scissors in two animals. We first recorded the activity of several motoneurons before and several after the transsection. To quantify the contribution of voltage dependent sodium channels to single motoneuron activity, QX 314-containing electrodes (100 mM in 2 M KCOOH) were used to impale motoneurons and QX314 was electrically driven into the neurons via current injections. The firing patterns directly after the impaling of the neuron and after QX314 injection were compared in the same neurons.

To block gap junction coupling, a carbenoxelone solution (CBX, 10 mM in 0.1 M PB) was injected into the VMN with micropipettes (tip diameters, 20–30 μm). Due to the long time course of gap junction blockage, CBX was also superfused over the fourth ventricle that lies directly above the VMN (see Fig. 1c) (Beaumont & Maccaferri, 2011; Rozental et al., 2001). After waiting 30-40 mins for the CBX to take effect, recordings were resumed.

Strychnine or bicuculline (10 mM in 0.1 M PB) was applied as described above for CBX to block glycine or GABA receptors, respectively. After baseline recordings, the pipette solution was pressure-ejected using a picospritzer (Biomedical) set to deliver three pulses, 10– 50 ms duration each, at 25–30 PSI at three locations along the rostro-caudal axis of the VMN. Baseline and post-injection vocal activity was recorded from five motoneurons prior to and after strychnine/bicuculline injections in multiple animals.

Lastly, 3 M KCl-filled electrodes were used to increase intracellular chloride concentration in order to reveal inhibitory input. After electrode penetration and a brief recording of baseline activity, current injection was used to drive chloride ions into the recorded motoneuron, after which its activity was further recorded.

### Data analysis

Neuronal data were analysed using Igor pro 6 (Wavemetrics), the software package neuromatic (http://www.neuromatic.thinkrandom.com) with custom written scripts (B.P.C.). The firing pattern of VMN neurons was visualized using a phase plane plot of the recorded voltage (*V*) against the difference in voltage over time (d*V*/d*t*). Statistical analysis was performed with the software Sigmaplot. A paired t-test was used to compare intracellular activity across neurons before and after pharmacological application using QX314 or KCl. We tested for a normal distribution and used a Mann Whitney U test where the normal distribution failed for comparison of activity across neurons before and after dorsal root cutting or application of bicuculline, strychnine, or CBX. Analysis of action potential and afterhyperpolarization amplitude evoked by antidromic stimulation were measured as follows: the baseline was evaluated by averaging 10 s prior to the antidromic stimulus; the maximum and minimum amplitude was then determined during a 30 s window after the antidromic stimulation pulse for multiple traces of different neurons. To account for differences in resting membrane potential across neurons, we calculated the voltage difference of the minimum and maximum from the baseline value expressed as %. The same analysis was performed during VOCs.

### Vocal motoneuron labeling and immunohistochemistry

Surgical and nerve labeling methods follow those in prior neuroanatomical studies (Bass et al., 1994; Chagnaud & Bass, 2014). As with neurophysiology (see above), animals were deeply anesthetized during all surgical procedures by immersion in aquarium water containing 0.025% benzocaine. Following ventral exposure of one vocal nerve at the rostral pole of the swim bladder, crystals of 10 kDa dextran-biotin (Molecular Probes, Eugene, OR, USA) were applied to the cut end of the nerve. Two juvenile males were deeply anesthetized by immersion in aquarium water containing 0.025% benzocaine and then transcardially perfused with 4% paraformaldehyde and 0.5% glutaraldehyde in 0.1 M phosphate buffer (PB; all Sigma); survival times were two (5.3 cm standard length) or four (10.8 cm) days. Brains were postfixed for 1 h, and then washed and stored in 0.1 M PB at 4° C. Prior to sectioning, brains were transferred to 30% sucrose in 0.1 M PB overnight at 4° C. Brains were embedded in Tissue-Tek O.C.T. compound (Sakura Finetek, Torrence, CA, USA), cryo-sectioned at 25 μm in the transverse plane, and collected onto Superfrost Plus slides (ThermoFisher Scientific, Waltham, MA, USA). Slides were stored at −80° C.

Slides were rehydrated in 0.1 M PB-saline (PBS, 2 x 15 min). To reduce background autofluorescence from glutaraldehyde, slides were incubated in freshly prepared 0.1% sodium borohydride in PBS for 10 min, followed by washes in PBS (3 x 5 min). Slides were blocked in 10% normal goat serum with 0.5% Triton 100 in PBS (PBS-NGS-T) for 2 h, then incubated overnight (18 hours) with the following antibodies: anti-glycine (1:200, rabbit, polyclonal, MoBiTec, 1015GE, Gottingen, Germany; see Rosner et al., 2018 for details on specificity and prior use in other vertebrates, including fish), anti-synaptic vesicle protein 2 (SV2, 1:50, mouse, monoclonal, Developmental Studies Hybridoma Bank, Iowa City, IA, USA). The SV2 antibody labels a transmembrane transporter in synaptic vesicles, recognizes all three known isoforms and has demonstrated specificity in both mammals and non-mammals (Buckley and Kelly 1985, manufacturer’s information). It labels synaptic vesicles in numerous teleost species (e.g., Buckley & Kelly, 1985; Schikorski et al., 1994; Smith et al., 2000; Stil & Drapeau, 2016). Slides were washed in PBS with 0.5% Triton 100 (PBS-T, 4 x 5 min), incubated 4 h with goat anti-rabbit AlexaFluor 568 (1:200), goat anti-mouse AlexaFluor 647 (1:200) and 488 anti-Streptavidin (1:500) in PBS-NGS-T, washed in PBS-T (4 x 5 min) and coverslipped using Vectashield (Vector Labs, Burlingame, CA, USA) and 1.5H coverglass (Thorlabs, Newton, NJ, USA).

### Super-resolution microscopy

Multi-color images using super-resolution structured illumination microscopy (SR-SIM) were acquired with a Zeiss Elyra S.1 SIM system using a 63x/1.4 oil immersion lens and ZEN 2012 software (Zeiss). Z-stacks (0.1 μm step-size, 3 μm thick) were acquired using five grid rotations and five phases at each plane. To maximize signal-to-noise, images were acquired within the first 5 μm from the tissue/coverglass interface. SR-SIM images were processed in ZEN, followed by generation of maximum projections and single optical plane images with ImageJ.

## Acknowledgements

Research support from NSF IOS 1457108 and 1656664 to A.H.B., DFG CRC 870/B17 to B.P.C., and NIH 1428922 to Cornell University Biotechnology Resource Center Imaging Facility (Zeiss Elyra super resolution microscope, SRM). We thank L. Remage-Healey for toadfish boatwhistle recording and Becky Williams for SRM guidance.

## Supplementary figures

**Supplementary Figure 1:**
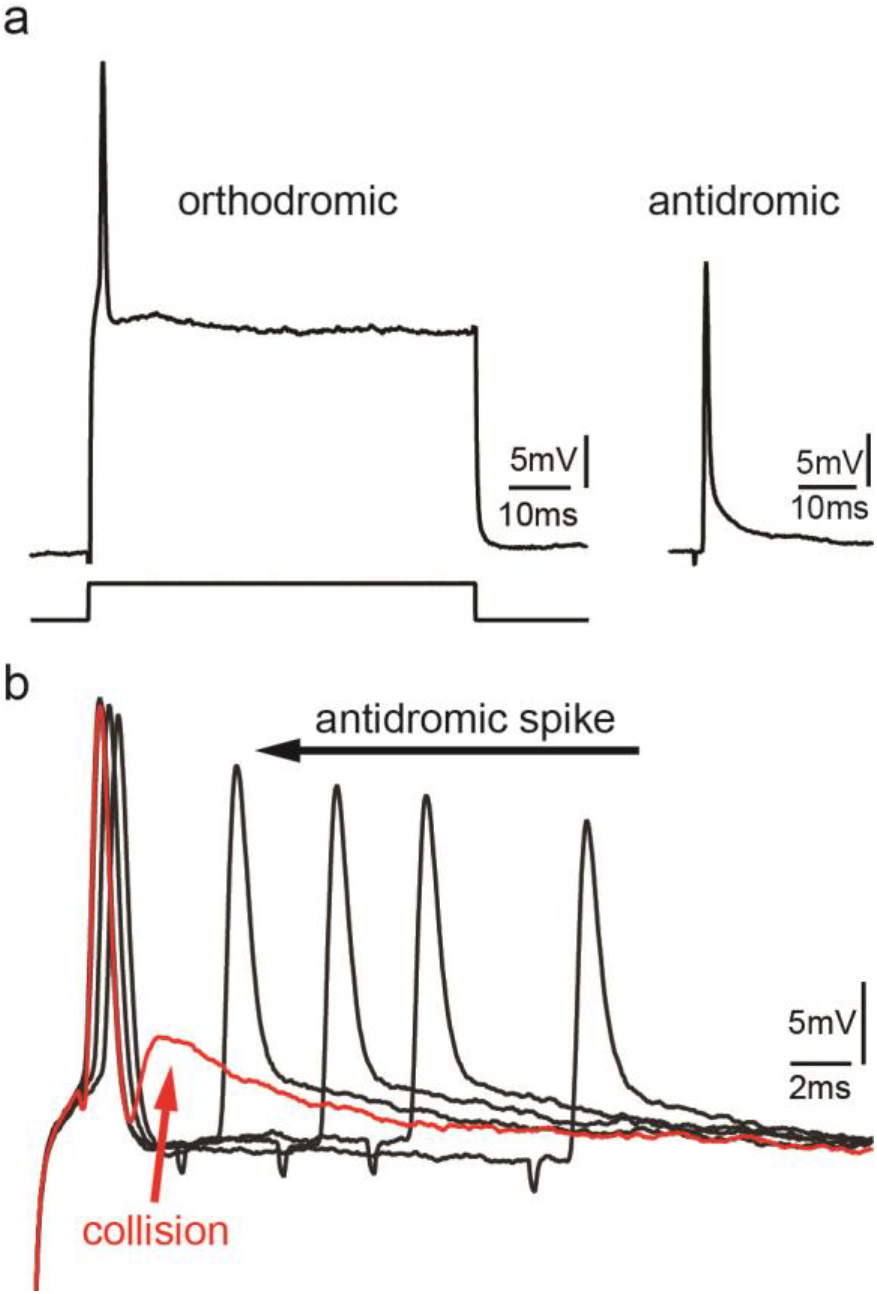
a) Action potential firing in a vocal motoneuron after intracellular current injection (left) and antidromic stimulation via the vocal nerve (right). b) Combination of intracellular current injection and antidromic stimulation at different latencies (black arrow) reveal a depolarizing potential when action potentials collide (red trace and arrow).

**Supplementary Figure 2:**
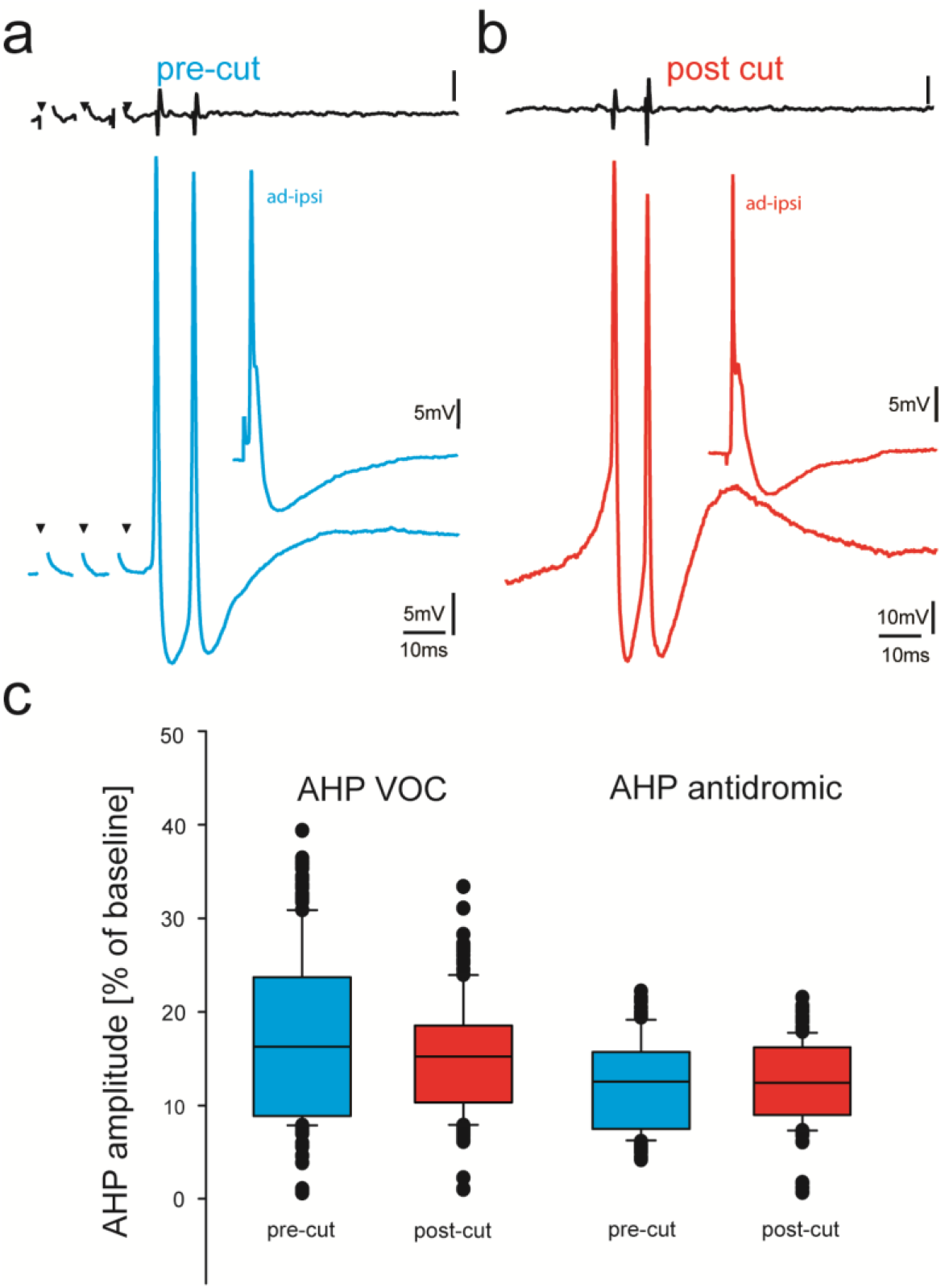
a) Midbrain evoked VOC recording from vocal nerve (top trace, black) and intracellular recording in a vocal motoneuron before (blue) and in another motoneuron in the same fish after transection of the dorsal root input to the hindbrain (red). Inset shows antidromic stimulation of the respective motoneurons. b) Box and whisker plots of after hyperpolarization (AHP) amplitude during vocal activity (VOC) and antidromic stimulation. Pre-cut (blue) and post-cut (red) conditions are indicated.

**Supplementary Figure 3:**
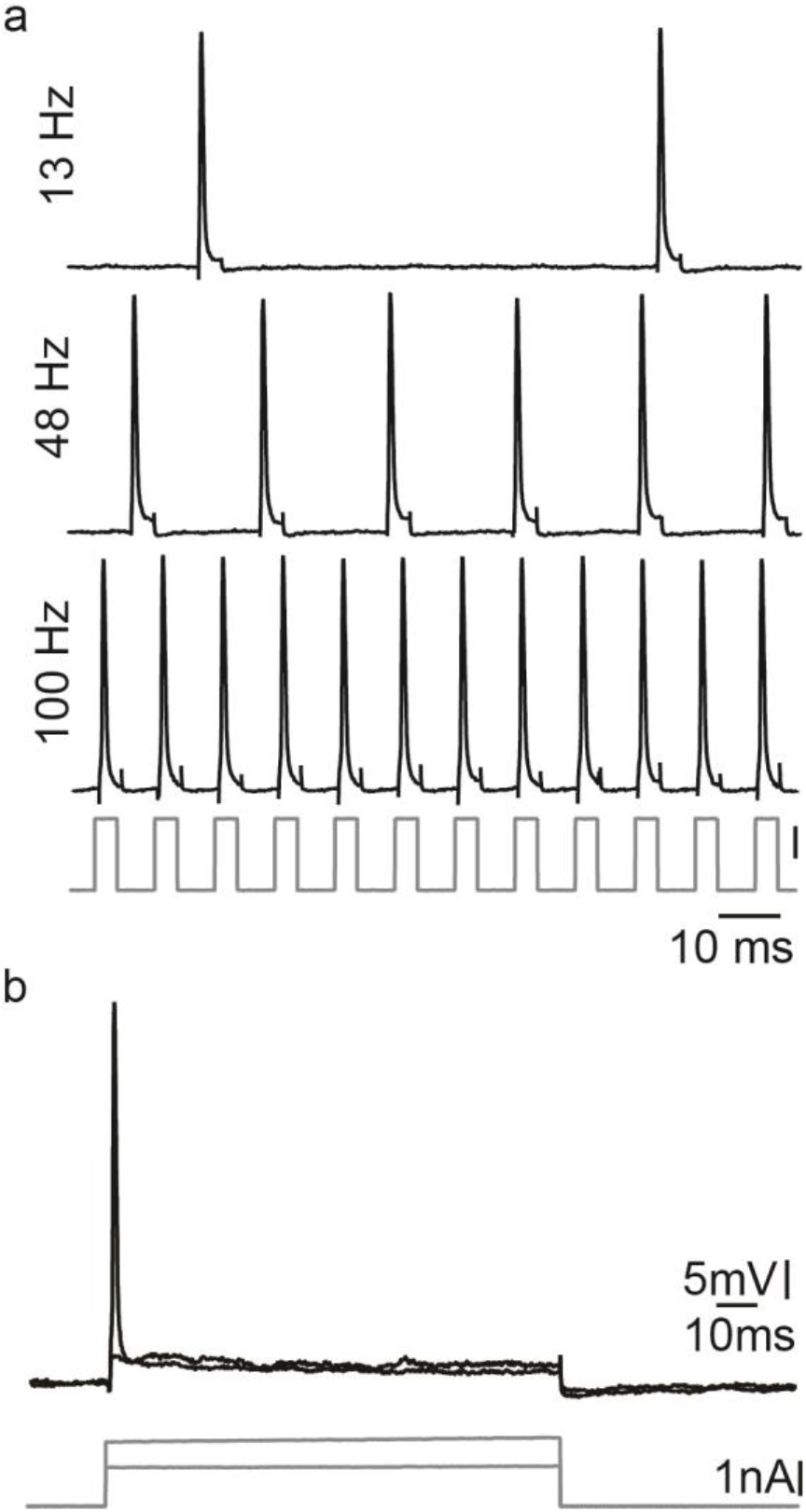
a) Action potential firing after short intracellular current pulse injection into a vocal motoneuron (at different stimulation frequencies). b) Intracellular long-lasting current injection into the same vocal motoneuron shows the necessity of repeated pulse stimulation to repetitively elicit action potentials in vocal motoneurons.

Supplementary movie 1: Intracellular recording of a vocal motorneuron during sequences of repeated antidromic stimulation via the vocal nerve on the ipsilateral and contralateral sides (both subthreshold to network activation of glycinergic input). The latency between the ipsi- and contralateral stimulation was diminished, which eventually led to the generation of an afterhyperpolarization (AHP) via glycinergic input.

Supplementary movie 2: Intracellular recording of a vocal motorneuron during sequences of repeated antidromic stimulation via the vocal nerve on the ipsilateral and contralateral sides. The latency between the ipsi- and contralateral stimulation was diminished, which eventually abolished the activation of the glycinergic input as motoneurons were hyperpolarized.

